# Contextual Expectations in the Real-World Modulate Low-Frequency Neural Oscillations

**DOI:** 10.1101/2024.05.30.596613

**Authors:** Victoria I. Nicholls, Alexandra Krugliak, Benjamin Alsbury-Nealy, Klaus Gramann, Alex Clarke

## Abstract

Objects in expected locations are recognised faster and more accurately than objects in incongruent environments. This congruency effect has a neural component, with increased activity for objects in incongruent environments. Studies have increasingly shown differences between neural processes in realistic environments and tasks, and neural processes in the laboratory. Here, we aimed to push the boundaries of traditional cognitive neuroscience by tracking the congruency effect for objects in real world environments, outside of the lab. We investigated how neural activity is modulated when objects are placed in real environments using augmented reality while recording mobile EEG. Participants approached, viewed, and rated how congruent they found the objects with the environment. We found significant differences in ERPs and higher theta-band power for objects in incongruent contexts than objects in congruent contexts. This demonstrates that real-world contexts impacts how objects are processed, and that mobile brain imaging and augmented reality are effective tools to study cognition in the wild.

## Introduction

Every day we explore our environment, look at, and recognise objects. Being able to recognise objects quickly is important, for example, when crossing a road it is important to recognise if a vehicle is approaching to prevent a collision. Specific objects are generally experienced in certain locations and environments, knowledge about which we learn through experience in the form of schemas. This knowledge can aid recognition, such that objects which appear in expected (or congruent) environments are typically recognised faster and more accurately than objects which appear in unexpected (or incongruent) environments (Bar, 2004; Biederman, Mezzanotte, & Rabinowitz, 1982; Brandman & Peelen, 2017; Cornelissen & Võ, 2017; Davenport & Potter, 2004; Henderson, Weeks, & Hollingsworth, 1999; Loftus & Mackworth, 1978). This congruency effect also has a neural component, with studies showing a larger neural response for incongruent objects between around 250 and 500 ms after the object appears, present in ERPs (e.g. the N400) and in theta oscillations (Coco, Aruajo, & Peterson, 2017; Coco, Nuthman, & Dimingen, 2020; Draschkow, et al, 2018; Bastiaansen, et al, 2005; Willems, Oostenveld, & Hagoort, 2008; Davidson & Indefry, 2007; Klimesch 1999). These findings have been linked to the integration of new semantic information into memory when the item was more unexpected (Bastiaansen, et al, 2005; Willems, Oostenveld, & Hagoort, 2008; Klimesch 1999; Riddle, et al, 2020; Metzner, et al, 2015; Hald, Bastiaansen, & Hagoort, 2006), although recent computational modelling work shows that N400 effects during language comprehension reflect a lexico-semantic prediction error signal (Eddine, et al., 2024). However, the majority of studies investigating the congruency effect have been conducted in well-controlled laboratory environments, where objects and scenes are presented together on a computer monitor before being replaced by an unrelated object and scene, breaking the temporal and spatial coherence that structure real-life experiences. Moreover, the participant is typically separated from the environment the objects are in, instead experiencing the ‘environment’ while being in a testing cubicle. This is different from how the real world is experienced, where there is a clear spatiotemporal coherence in the environment. As a consequence, we have little information about whether the tasks participants perform in laboratory settings are a good approximation of behaviours and associated neural processes in the real world. Indeed, past research indicates differences between neural processes in realistic environments and tasks, and neural processes in the laboratory (David, Vinje, & Gallant, 2004; Aghajan, et al, 2015; Hölle, & Bleichner, 2023; Hampton, Hampstead, & Murray, 2004; Redcay, & Schillbach, 2019; Maimon, Straw, & Dickinson, 2010). This raises the question to what extent do research findings translate to real environments, and stresses the importance of pursuing neurocognitive issues within their true dynamic functional context.

There is increasing acknowledgment that neuroscience must seek out more naturalistic ways to study cognition (Nastase, Goldstein, & Hasson, 2020; Gramann, et al, 2014; Grasso-Cladera, et al, 2024; Shamay-Tsoory, & Mendelsohn, 2019; Stangl, Maoz, & Suthana, 2023), however significant challenges remain for taking our experimental techniques out of the lab. Developments in mobile electroencephalography (mEEG) technology over the last ten to fifteen years, such as the development of the MoBI framework (Makeig, et al, 2009) and improved signal processing tools (Klug, et al, 2022), mean that mEEG has become a reliable method which provides the opportunity to perform neuroimaging in the real world (Gramann, et al, 2014; Makeig, et al, 2009; Niso, et al, 2023; Griffiths et al, 2016; Mustile, et al, 2023; Ladouce, et al, 2019). However, in experiments, we typically want to control or manipulate some aspect of what we study, which poses technical barriers when experiments are situated in the real world. The development of mixed reality methods, such as head-mounted augmented reality (AR), allows researchers to control when and where stimuli are presented to participants while they freely move around the real world. While AR has been utilized to study assembly systems (Wang, Ong, & Nee, 2016), pedagogy (Saidin, Halim, & Yahaya, 2015) and treatment of psychological disorders (Chicchi Giglioli, et al, 2015), a relatively novel approach is to use mobile EEG and head-mounted AR together, to study the neural processes underpinning cognition in real settings (Krugliak, & Clarke, 2022).

Here we study how neural processes are modulated by the relationship between the object and the environment, by presenting virtual objects in real locations using AR, while neural activity is recorded with mobile EEG. Participants walked around a large indoor and outdoor area seeing virtual objects that were either more or less congruent with their setting. Concurrent EEG then allows us to ask how the processing of objects is modulated by the congruency between the objects and the environment, thereby allowing us to test if our knowledge about the world modulates the processing of visual objects.

## Materials and Methods

### Overview

The initial challenges were first to create a mixed reality experiment where objects were embedded into the real environment, set to appear at specific locations when the participant was in proximity to an augmented marker, and second, to have sufficient synchronisation between the AR and EEG systems for millisecond precision for onset markers of augmented objects. We used a large indoor and outdoor space near the Department of Psychology at the University of Cambridge as our testing arena, with the 80 virtual objects being assigned a physical location (Fig. 1C). The objects and locations were chosen such that 40 objects would be more congruent and 40 less congruent with the environment.

**Fig. 1.**
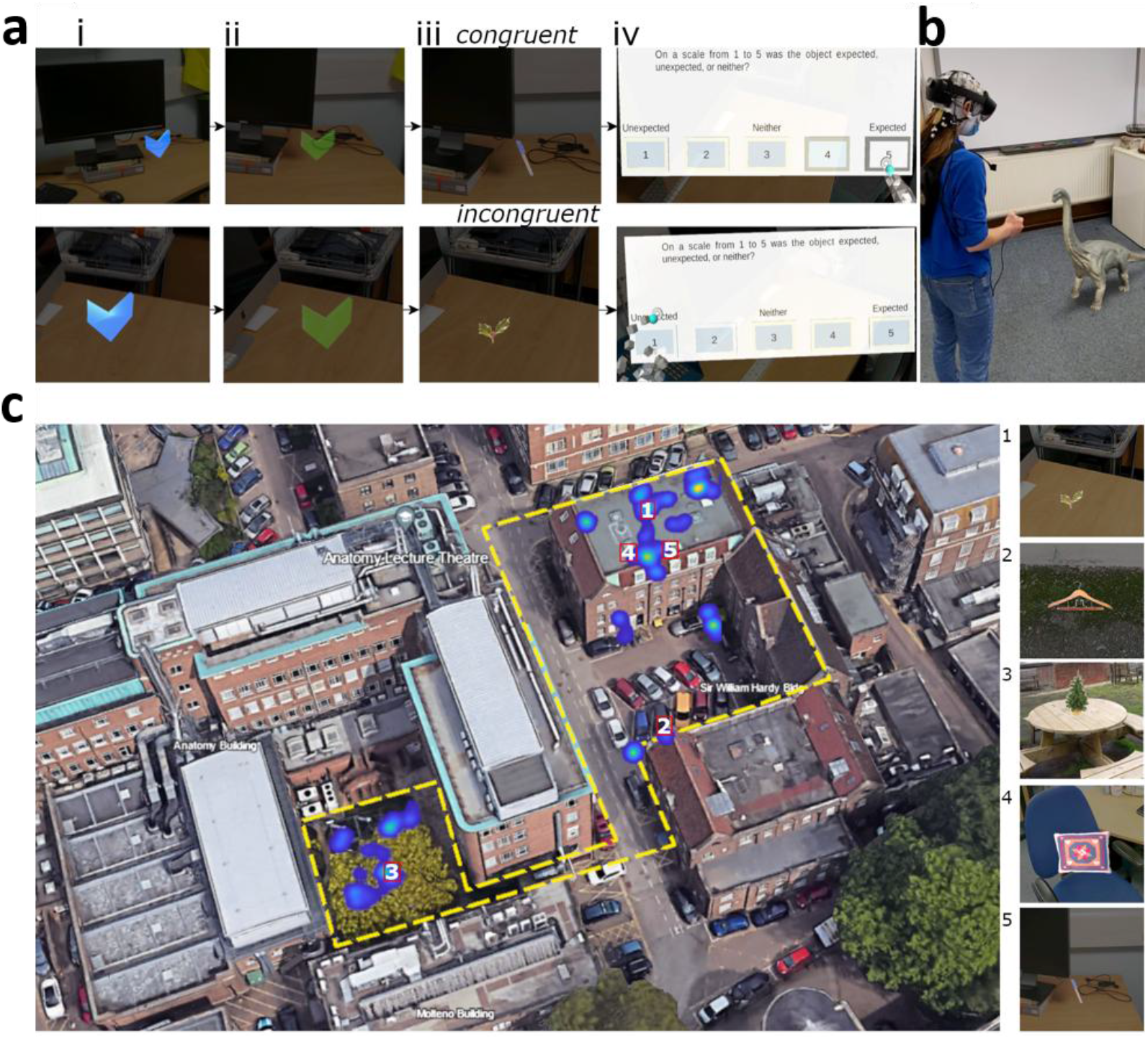
Experimental paradigm, example setup, and object locations. (Ai). Example of the blue virtual arrow indicators participants searched for, these indicated the object locations. (Aii) The blue arrow turns green when participants moved close enough to the arrow. (Aiii) Example object which appeared for 5s after participants pressed the button on the button box once the arrows had turned green. (Aiv) Participants were presented with a congruency rating question once the object disappeared. For panel (A) the top row shows an example congruent trial, and the bottom row an incongruent trial. (B) Equipment setup illustrated on one of the authors who consented to having their picture taken and shared. The author is wearing the Brain Products LiveAmp EEG system with the Hololens 2 placed on top of the EEG system. The author is holding the button box that was connected to the Hololens 2 and the LiveAmp. An example virtual object (Brachiosaurus) is presented to the author. Note, a Brachiosaurus is not one of the objects presented to the participants. (C). A birds-eye view of the experimental environment taken from Google Earth v.10.55.01 (May 15^th^ 2024), Cambridge, UK, and edited. The yellow dashed line indicates the full area that objects were placed within. The heatmap indicates locations a participant gazed at the objects, and example objects in locations 1-5 are presented on the right-hand side. Objects are ordered by the predefined congruency ratings.

Once the participants were fitted with the mobile EEG system, the AR device was placed over the top of the electrodes. We used an initial calibration to align the real and virtual worlds, after which the experiment began. On each trial, a virtual blue arrow appeared at the location of the object (Fig.1.Ai). To facilitate finding the object, the arrow could be seen through walls and surfaces. Once the participant was near the arrow, it turned green (Fig.1.Aii), signalling the participant could press a button after which the arrow would change into an object (Fig.1.Aiii). The object was presented for 5 seconds, after which, a virtual display appeared to collect an expectancy rating on a 5-point scale, which is used to accommodate individual differences in how congruent each item was with its setting (Fig. 1.Aiv). The experiment had four blocks, two indoors and two outdoors, with each block containing 20 objects, 10 of which would be more congruent and 10 less congruent. The order of the blocks was counter-balanced between participants. The study followed the procedures described in our previous protocol article (Nicholls, et al, 2023), and an additional exploratory analysis of the time-frequency data.

### Participants

To determine the number of participants required, a power analysis was conducted on data from Draschkow, et al, (2018) who examined EEG congruency effects of object-scene processing. The power analysis was conducted using the ‘sampsizepwr()’ function in MATLAB (MATLAB, 2021), based on the mean difference between the amplitude of the N400 for congruent and incongruent conditions, to obtain a power of 0.8, at an alpha of 0.05. The analysis showed that a minimum sample size of 34 participants was required.

Overall forty-two participants took part in the experiment (13 female, aged 18-35 years). Eight participants had exceptionally noisy data and were excluded from the analysis, leaving a total of 34 participants for the final analyses in line with the outcome of the power analysis. All participants had normal or corrected to normal vision and reported no history of neurological conditions. Ethical approval for the study was granted by the ethics committee of the Department of Psychology at the University of Cambridge. Participants signed informed consent prior to the start of the experiment. This study was performed in accordance with all appropriate institutional guidelines and international guidelines and regulations, in line with the principles of the Helsinki Declaration.

### Apparatus

Participants were presented with virtual stimuli using a Hololens 2 device (https://www.microsoft.com/en-us/hololens/). The Hololens 2 is a mixed reality headset with a horizontal field of view of 42°, a vertical field of view of 29°, and presents images at 60 Hz. EEG was recorded using the Brainvision LiveAmp 64 mobile system (Brain Products GmbH, Gilching, Germany), with an online low pass filter of 131 Hz. We recorded 64-channel EEG through ActiCap Slim active Ag/AgCl electrodes located in an elastic cap (actiCAP snap) positioned according to the international 10/20 system (Jasper, 1958), with a reference electrode placed at FCz, ground electrode placed at Fpz, and a sampling rate of 500 Hz. The EEG electrodes were placed on the participant with the Hololens 2 placed on top of the electrodes, and the LiveAmp and electrode cables placed in a backpack that the participant wore while performing the experiment. A custom-built button box was plugged into both the Hololens 2 USB-C port and the LiveAmp 1-bit trigger port, meaning that when the button was pressed, it simultaneously sent a signal to Hololens 2 and a 5V signal to the LiveAmp. The LiveAmp then converted that signal to a trigger that was marked in the EEG recording. The signal to the Hololens 2 triggered the appearance of an object during the experiment. An example of the setup can be seen in Fig. 1.B.

### Stimuli and environments

During the experiment, participants explored a large real-world environment, including indoor (consisting of three offices, a corridor, and a meeting room) and outdoor elements (consisting of a parking lot and a picnic area) where they were presented with virtual objects in various locations (Fig. 1.C). A total of 80 images corresponding to concepts from a property norming study (Hovhannisyan, et al, 2021) were presented to the participants, and five additional images used as practice trials.

The 80 images used in the main experiment were selected from the one thousand images in the property norms set. The one thousand images were narrowed down to 400 by the lead author, based on which objects could plausibly appear in the experimental location. These objects were then rated as either congruent or incongruent with the indoor and outdoor environments. From this we extracted 40 objects that were the most expected in the environments, and 40 objects that were the most unexpected in the environments. For the unexpected objects we made sure that they were highly unexpected, but not impossible. For example, a reindeer in a parking lot in the UK is highly unexpected but not impossible. However, if an object would be highly unexpected and impossible, we would not use that object. The congruency ratings for the 80 objects were then verified by a second experimenter. All objects were placed in plausible locations within the environments (i.e. a bottle on a table, not floating in the air). These congruency ratings were used to form the congruency conditions in one of our analyses.

### Constructing the paradigm

To place the stimuli in the correct locations in the real world, a virtual 3D map of the real-world testing area was created. This involved scanning the experimental environment through the Hololens 2, which uses spatial mapping to create detailed representations of real-world surfaces which are built into a map. This spatial map was then downloaded from the device, and imported into Unity for use as a virtual environment to place the objects within. The locations used in the virtual environment would correspond to the matching locations in the real world (an example is shown in Fig. 2). The experiment was coded using SilicoStudio (beta testing version, https://www.silicolabs.ca/), an experimental design software package built for the Unity engine, and the experiment was presented to participants through the Hololens 2. SilicoStudio was used to place the objects in the correct real-world locations and to enable interactions between the participant and the objects. The locations of the stimuli were the same for all participants. A list of the stimuli used and their locations in the experiment can be found in the Supplementary Materials (Supplementary Table S1, Supplementary Figs. S1 and S2). The interactions included the changing of the arrow colour from blue to green when participants stepped close enough. The second interaction included changing the indicator into an object when the participants pressed the button on the button box. Furthermore, SilicoStudio was used to manage the order in which objects appeared, and to record behavioural data which included head and eye tracking (not analysed here), and behavioural responses.

**Fig. 2.**
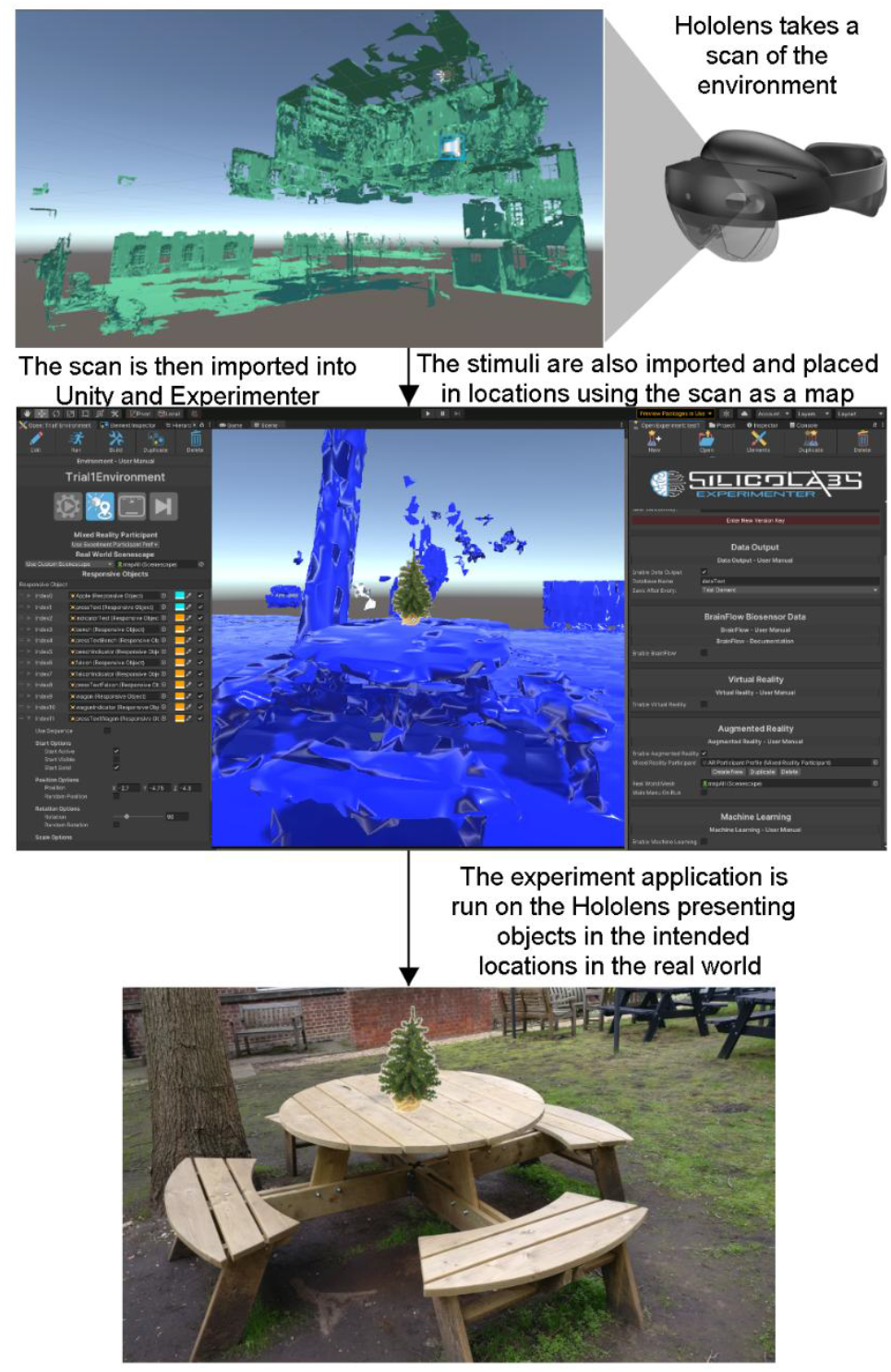
A description of how the experiment was created. First the Hololens 2 is used to take a scan of the real world. This is done by walking around the real world and looking at as many parts of the world as possible. For example, to scan a door one would first look at the top left corner of the door then move their head down slowly to the bottom left corner. Then look slightly to the right and move your head slowly upwards. This would be repeated until you had looked at all parts of the door. As you are looking and scanning the environment, the Hololens constructs 3D models or maps of the real world, an example of our environment is shown in A. This 3D map can be downloaded from the device and then imported into Unity, shown in panel B, and used in the same way as a virtual environment. The stimuli were then placed in the intended locations. As the 3D map is an accurate map of the real world, the locations the objects are placed in within Unity correspond to their real world locations. Therefore, when exporting the virtual environment, with the stimuli objects, back to the Hololens 2, the device will present the objects in the intended locations in the real world (As in panel C).

### Procedure

The 80 images were presented to participants in four blocks of 20. The images in each block were matched on the following predictors: congruency, category, domain, hit rate on a visual task, hit rate on a lexical task, false alarm rate for a lexical task, false alarm rate for a visual task, image energy, image size, proportion of non-white space in the image, hue of the image, saturation of the image, frequency in the COCA database, proportion of participants identifying the modal name of the object, and the number of non-taxonomic features. Except for congruency and environment, the predictors for each image were taken from data included with the property norming data related to our stimuli (Hovhannisyan, et al, 2021). The matching was performed using the Match software (version 2.09.02, van Casteren & Davis 2007).

To familiarise the participants with the Hololens and the appearance of AR stimuli, they performed five practice trials in an indoor space. The trials followed the same procedure as for the experiment described below. During the experiment participants walked through an environment, with indoor and outdoor areas. Participants completed two blocks of 20 trials indoors, and two blocks of 20 trials outdoors. Each block of 20 trials contained 10 that were more congruent and 10 that were more incongruent. In each trial participants needed to find an arrow indicating the location of the object (Fig. 1.Ai). When participants were close enough to the arrow, the arrow changed colour (Fig. 1.Aii) indicating that the participants could press the button on the button box (Fig. 1.A). Once the button was pressed, an object appeared for five seconds (Fig. 1.Aiii). At the same time a trigger was sent to the 1-bit TTL input of the LiveAmp by the button press. Participants were instructed to look at the object for the entire time it was visible, and to keep as still as possible. After the object disappeared, participants were presented with a virtual screen, asking them how expected or congruent they found the object within the environment on a scale from one to five (Fig. 1.Aiv). One was unexpected (incongruent), three was neither, and five was expected (congruent). Participants indicated their rating by pressing the appropriate button on the virtual screen. These ratings are used for the participant rating analysis. We included these ratings as participants may have different ideas or experiences of which objects are congruent and incongruent in the environments. The next trial began once they had responded. This process was repeated until all 80 objects were found. During the whole session, the participants’ EEG activity were recorded. The four blocks took approximately 60 minutes to complete.

### EEG Pre-processing

Pre-processing was performed using the EEGLAB toolbox (Delorme & Makeig, 2004) with the BeMoBIL pipeline (Klug, et al, 2022), which allowed for the semi-automated analyses of mobile brain imaging signals. The single-subject data were truncated by removing parts of the EEG recording that occurred before the start of the experiment, the inter-block intervals, and after the end of the experiment. The data were then downsampled to 250 Hz and power line noise was cleaned using the Zapline toolbox (de Cheveigné, 2020). Noisy channels were automatically identified using the EEGLAB clean_artifacts() function with the maximum tolerated fraction of the recording duration during which the channel was flagged as ‘bad’ was set to 0.5 and the minimum channel correlation was set to 0.75. Otherwise the template configuration settings from the BeMoBIL pipeline were used for the clean_artifacts() function. The identified bad channels were then interpolated (mean interpolated = 3, SD = 3) and an average reference was applied to the data.

At this point, the dataset was duplicated so that IC weights could be computed on a filtered version of the data and then applied to a non-filtered dataset. For ICA only, the duplicated data were highpass filtered using a zero-phase Hamming window FIR filter with a cutoff frequency of 1.5 Hz (order 1650, transition width 0.5 Hz). Independent components (ICs) were then identified using an adaptive mixture component analysis (AMICA; Palmer, et al, 2008). The AMICA decomposition uses an automatic time domain cleaning where samples are removed based on the log likelihood that the samples do not correspond to the algorithm’s estimate of model fit. Three standard deviations were used as the removal criterion. The ICs identified by AMICA were copied into the original unfiltered dataset, and classified as ‘brain’, ‘eye’, ‘heart’, ‘channel noise’, ‘line noise’, ‘muscle’, and ‘other’ using IClabel Lite (Pion-Tonachini, Kreutz-Delgado, & Makeig, 2019). ICs that were labelled as ‘eye’, ‘heart’, ‘channel noise’, and ‘line noise’ were then excluded from the reconstructed dataset (mean excluded = 8, SD = 3). Another version of the data analysis where ICs labelled as ‘muscle’ were also removed are shown in Supplementary Fig. 3 in the Supplementary Materials. All effects reported in the Results remained when ‘muscle’ components were additionally removed. However, as IClabel is trained on a very small portion of mobile EEG data we are not confident that IClabel accurately labels ‘muscle’ ICs over ‘brain’ ICs (Klug, et al, 2022), therefore, we went with the preprocessing pipeline where we were confident no ‘brain’ ICs were removed.

The IC-cleaned data were then filtered with a one-pass zero-phase Blackman-windowed sinc FIR filter with cutoff frequencies of 1 and 40 Hz, and a transition width of 0.1 Hz for the ERP analyses, and cutoff frequencies of 1 and 100 Hz for the time-frequency analyses.

The data were then epoched between -2 and 3 seconds around the onset of the images. Channel baseline means were removed between -100 and 0 ms before the onset of the stimulus. Noisy epochs were automatically removed using the bemobil_reject_epochs() function, based on the mean of channel means (to detect general large shifts), the mean of channel standard deviations (SDs; to detect large fluctuations in many channels, e.g. large muscle activity), the SD of channel means (to detect phases where single channels had large fluctuations), and the SD of channel SDs (to detect large fluctuations in single channels). We used an automatic kneepoint with an offset of 5% to determine a threshold for epoch rejection. On average, 8 trials were rejected per participant (SD=4).

### EEG Analyses

We analysed event-related potentials (ERPs) and time-frequency representations (TFRs) from channel data using cluster-based permutation tests from the FieldTrip toolbox (Oostenveld, et al, 2011). We performed two complementary sets of analyses where our congruency levels were either defined using the predefined congruency conditions (40 congruent trials and 40 incongruent trials, 2 conditions), or where congruency was defined using the participants own ratings on the 5-point scale. The data from the indoor and outdoor blocks were pooled together so that our congruency effects reflect a wider range of environments and increase trial numbers per condition.

For the ERP analyses of the two congruency conditions, data were averaged across trials within the conditions giving two mean ERPs per channel per participant. For the ERP analysis we first attempted to replicate the data analysis approach taken in (Draschkow, et al, 2018). We computed the mean EEG amplitude in two response windows: between 250 and 350 ms (N300), and between 350 and 500 ms (N400), averaged across the central electrodes FC1, CP1, CP2, Cz, FC2, C1, CPz, and C2. We computed t-tests in the N300 and N400 time-windows between the congruent and incongruent conditions. As a more exploratory analysis, we also ran cluster-based permutation analysis across electrodes for data averaged within the N300 and N400 time-windows. A dependent samples t-test was performed at each electrode with a threshold of p = 0.025 to define clusters in the observed data using the maxsum calculation. A cluster-based permutation routine using 5000 randomisations was used to calculate cluster p-values for the observed data where an alpha of 0.05 was applied.

For the ERP analysis using the 5-levels from the participant ratings, we created 5 ERPs per participant based on their individual responses which had broadly similar numbers of trials (mean number of trials per condition; 1 (very incongruent) = 13.7 trials, 2 = 13.5 trials, 3 (neutral) = 11.3 trials, 4 = 14.8 trials, 5 (very congruent) = 14.4 trials). We tested for linear effects of congruency in the EEG data using the regression routine in Fieldtrip (depsamplesregrT) in conjunction with a cluster-based permutation test with the same parameters as above.

For the TFR analyses, we followed a similar process as for the ERPs. The TFRs were calculated for each trial using 5-cycle Morlet wavelets at 40 logarithmically scaled frequencies from 2 to 100 Hz and between -2 to 3 seconds around the appearance of the object. No baseline correction was applied in the calculations of the TFRs. The TFRs were calculated for each electrode individually resulting in two TFRs per electrode per participant for the congruency conditions analysis, and five TFRs per electrode per participant for the participant ratings analysis.

Cluster based permutation tests were then performed on the TFR data. For each electrode, the effect of congruency was tested between 2 and 100 Hz and between 0 and 1 second, using a cluster-defining threshold of 0.025 and 5000 randomisations. When there were two congruency conditions, congruency effects were assessed with a dependent samples t-test at each data-point. For the participant ratings analysis, a linear regression was performed determining the relationship between EEG amplitude and the 5 congruency rating, producing t-values for each electrode, time-point and frequency.

In all analyses, the significance of each cluster was determined by using the Monte Carlo method. For this the trials from all congruency level were placed in a single set. Trials are randomly drawn out from this set that match the number of trials for each congruency rating. The same analyses as above were then performed on the randomised dataset, and repeated 5000 times. A histogram was constructed from cluster masses of the randomised samples, and if the original cluster mass (from the observed data) was below the 2.5th or above the 97.5th quantile of the constructed histogram, they were considered to be significant at an alpha level of 0.05.

## Results

As a first step towards characterising the object recognition process in real world contexts, we examined the averaged EEG signals from the onset of the objects. Presenting participants with AR stimuli produced a clear visual evoked response over posterior electrodes, including a P100 followed by a negative deflection between 150 and 200 ms, and P300 components (Fig. 3a), with ERPs declining after approximately 600 ms. These peaks and characteristics were also present in the global field power of the EEG signals (Fig. 3b). A grand average time-frequency representation was also calculated (Fig. 3c), which showed a post-stimulus increase in low frequency power (lower-alpha/high-theta) between 100 and 400 ms, followed by a decrease in power between 500 and 1000 ms. There was a further increase in delta/theta power between 500 ms and 1000 ms, along with decreased gamma power throughout the post-stimulus period. These signals are in line with lab-based studies when presenting visual stimuli, suggesting that mobile EEG in combination with AR is a feasible approach to examining cognitive processes in real environments.

**Fig. 3.**
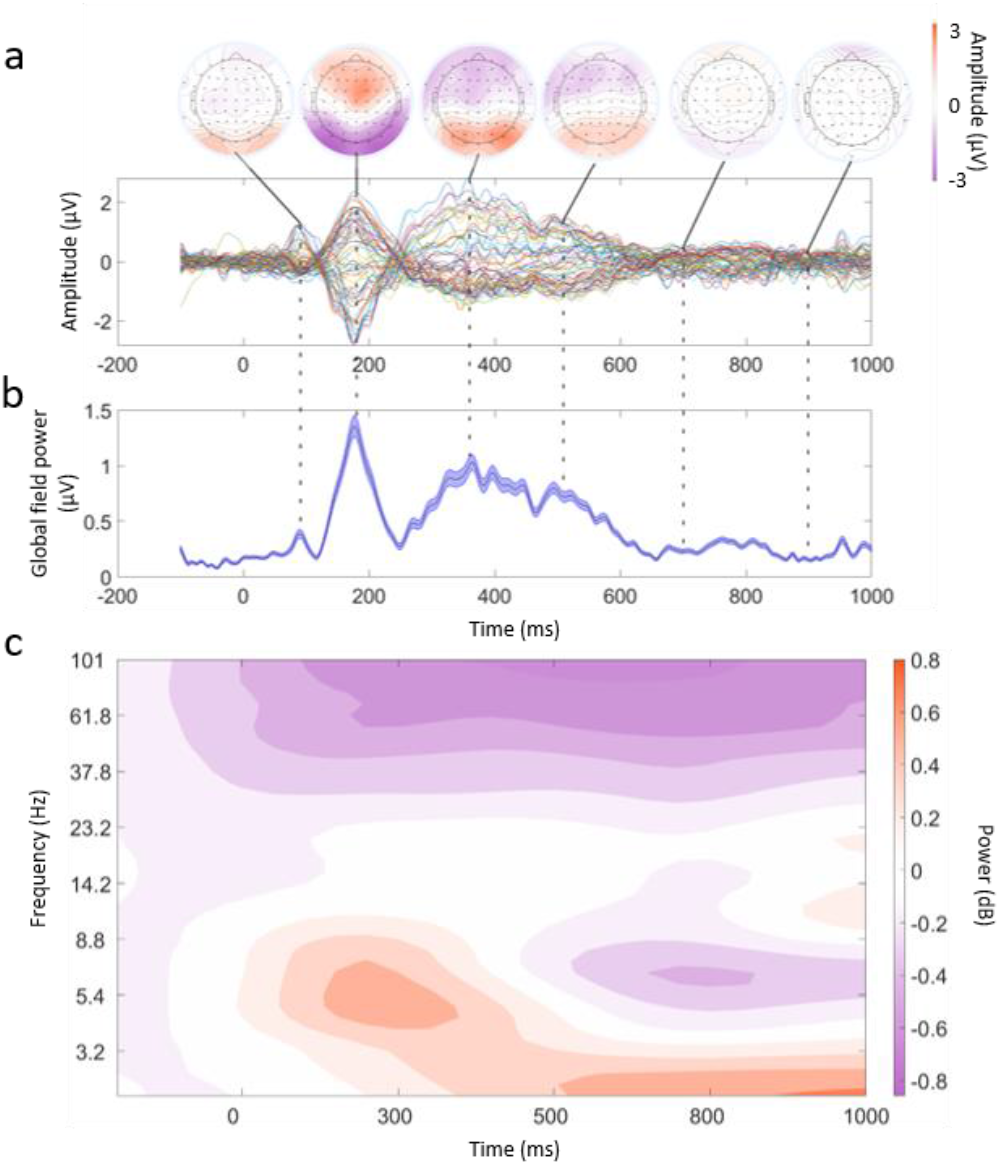
Example of mEEG data quality. (A) Butterfly plot averaged across all participants and trials. Each line corresponds to an EEG channel. Above this are topographical plots showing the average amplitude at timepoints of: 100, 170, 350, 500, 700, and 900 ms. (B) Global field power plot taken from the data used in A. The central line is the mean GFP and the lines above and below indicate the standard error. (C) Time-frequency representation of the EEG data from -100 to 1000 ms and frequencies 2-100 Hz. The frequencies are logarithmically scaled. The data was smoothed using a Gaussian kernel (3 Hz, 40 ms) across times and frequencies, and plotted using a filled 2D contour plot. A baseline correction was applied between - 100ms and 0ms for illustration purposes only.

We used two complementary approaches to determine if EEG activity was modulated based on the congruency between the objects and the environment. First, we defined object-environment congruency based on our predefined congruent and incongruent conditions which were matched on various factors and align with previous approaches. Second, we defined congruency based on the participants’ ratings given during the experiment on a 5-point scale. This second approach allows us to both account for individual variability in what people regard as congruous and examine congruency on a finer scale. These analyses will focus on both modulations of ERPs and of time-frequency modulations.

### Predefined congruency conditions

Previous lab-based EEG studies have typically shown a congruency effect in the N300 and N400 time range (250 to 500 ms) across central electrodes, with a larger negativity for objects that are incongruent with the environment compared to objects that are congruent with the environment. We attempted to replicate this finding using the same data analysis approach as in Draschkow et al (2018), and set out in our protocol paper (Nicholls et al., 2023). We computed the mean EEG amplitude in two response windows: between 250 and 350 ms (N300), and between 350 and 500 ms (N400), across the central electrodes FC1, CP1, CP2, Cz, FC2, C1, CPz, and C2. We found no significant effect of congruency in either time window (N300: t(33)= 0.07, SE=0.96, p=0.943; N400: t(33)=1.33, SE=1.02, p=0.194). Next, we expanded the analysis to test across all channels, which revealed a significant effect of congruency in the N400 time window over left anterior electrodes (cluster: AF7, F7, FT7; p = 0.025; Fig. 4a, b) and posterior central electrodes (cluster: Pz, O1, POz; p = 0.019; Fig. 4a, c). In both clusters, the incongruent condition showed a larger deviation from zero compared to the congruent condition, with the left anterior cluster displaying the anticipated modulation of an N400 response.

**Fig 4.**
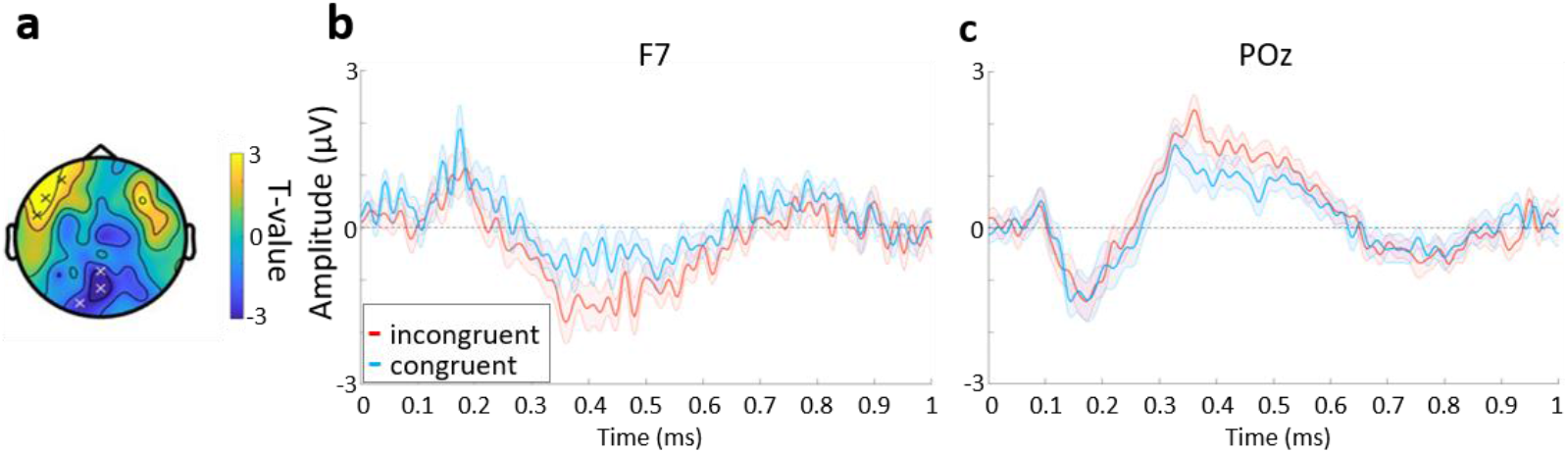
Congruency effects in ERPs. (a) Topographic plot of the statistical contrast between congruent and incongruent trials within the N400 time window. Electrodes in the significant clusters marked with an ‘x’ (frontal cluster: AF7, F7, FT7; posterior cluster: Pz, O1, POz). (b) ERP for electrode F7 from the frontal cluster, and (c) POz from the posterior cluster. In both (b) and (c)the blue line is the mean for the congruent trials, the red the incongruent trials, and the shaded area indicates the standard error.

Given the more fluid dynamics of real-world experiences, and that the context was ever present, we explored analyses less dependent on strict time-locking across trials to determine whether there were additional impacts of context on neural processing. As such, we explored whether the congruency between the object and the environment modulated different neural oscillations (Fig. 5). EEG power for each frequency and time-point was calculated for each trial, and averaged within each congruency conditions.

**Fig. 5.**
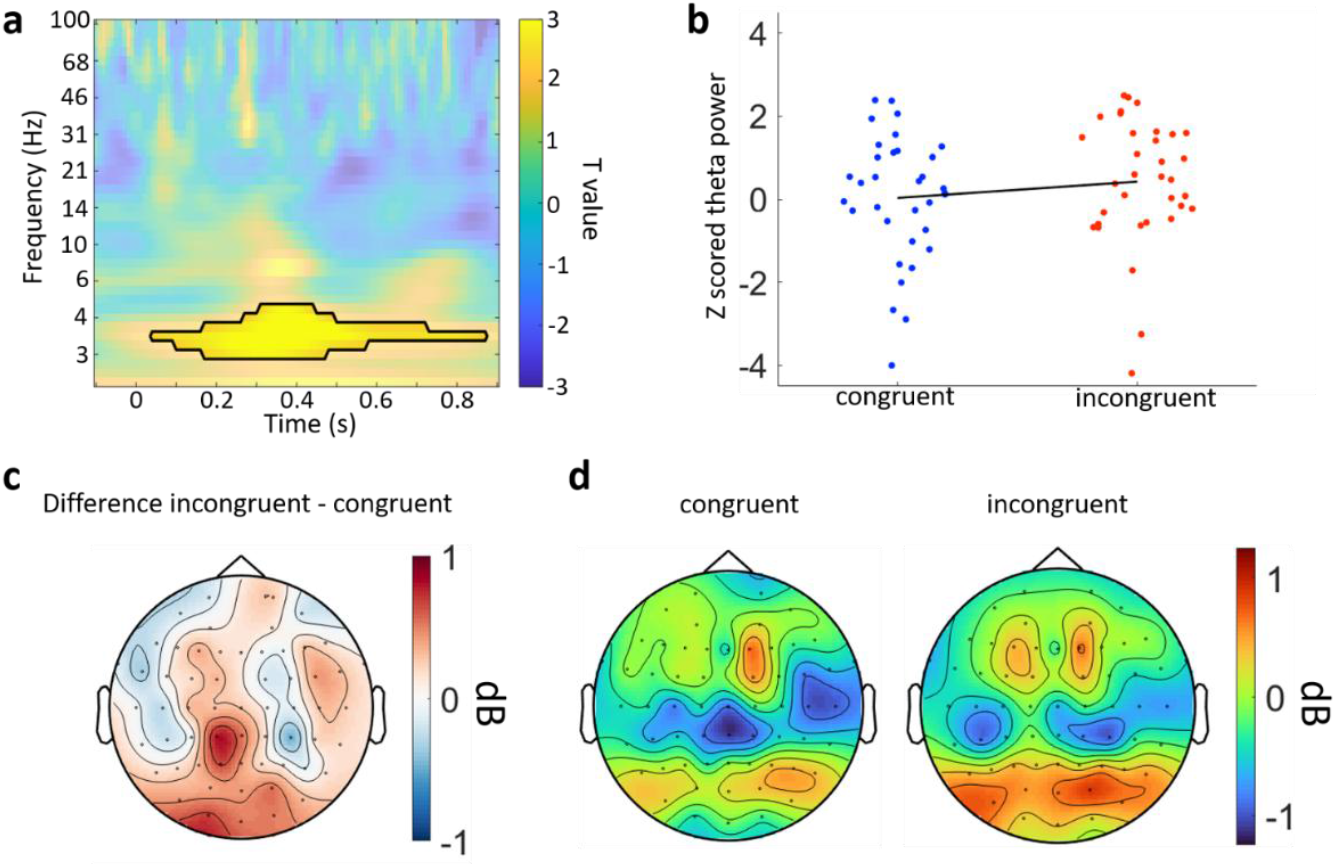
Time-frequency analyses of the congruency conditions. (a) Time frequency representation of the contrast between incongruent and congruent trials for electrode O1 from the posterior cluster. The black line indicates the significant cluster at this electrode. (b) Z-scored power from the significant cluster for each congruency condition. Each point is an individual participant. The black line is a first-degree polynomial fitted to the z-scored power. (c) Topography of the mean difference between congruency conditions averaged over 3 to 5 Hz power and between 200 and 600 ms. (d) Topographies for congruent and incongruent conditions showing the mean power averaged over 3 to 5 Hz and between 200 and 600 ms.

Our initial analysis targeted the two clusters where N400 ERP effects were seen. We ran cluster-based permutation analyses across time and frequency for the two clusters previously identified. This revealed a significant cluster where theta power was greater for the incongruent condition compared to the congruent condition which was centred between approximately 250 and 600 ms post-stimulus onset (Fig. 5a-d; 2-5 Hz; cluster p = 0.010). No additional effects were observed in the left anterior cluster, or when the analysis was expanded to test across all electrodes, time-points and frequencies. This shows that when more unexpected events occur, low-frequency neural responses increase over sensory and parietal electrodes in proportion to how unexpected the item was (Fig. 5a-d).

### Participant congruency ratings

We next explored how EEG activity was modulated by congruency using the ratings participants provided (Fig. 1a). This was because the participants’ experiences of what objects appear in which locations may differ, resulting in individual variability in which objects are considered congruent or incongruent with the environment. To capture this, and gain a more fine-grained understanding of the impact of congruency on neural processes, we asked participants to rate how expected (congruent) or unexpected (incongruent) the objects were using a five-point scale on each trial. Using these ratings, we can examine graded effects of congruency using linear regression-based analyses.

A linear regression was performed at each electrode and time point to determine if ERPs varied with congruency ratings. A cluster-based permutation test across time and electrodes found no evidence for a linear modulation of ERPs by the degree of congruency between objects and the real-world setting. This was also the case when our analyses targeted the N300 or N400 time-windows.

Next we explored whether the degree of congruency between the object and the environment modulated different neural oscillations (Fig. 6). EEG power for each frequency and time-point was calculated for each trial, and averaged within each of the 5 congruency ratings. We then ran a cluster-based permutation linear regression analysis across all electrodes, timepoints, and frequencies. This revealed a significant cluster showing a positive relationship where power increased with increasing incongruency between approximately 100 and 600 ms post-stimulus onset, centred in the theta range (Fig. 6a; 4-7 Hz; cluster p = 0.014). The cluster was primarily located over posterior electrodes, both centrally and on the right. This positive relationship indicates that there was significantly more power in the theta range between 100-600 ms for objects that were more incongruent with the environment compared to those that were more congruent. This shows that the more incongruous the object is with the environment, the greater the increase in low-frequency neural responses increase over sensory and parietal electrodes.

**Fig. 6.**
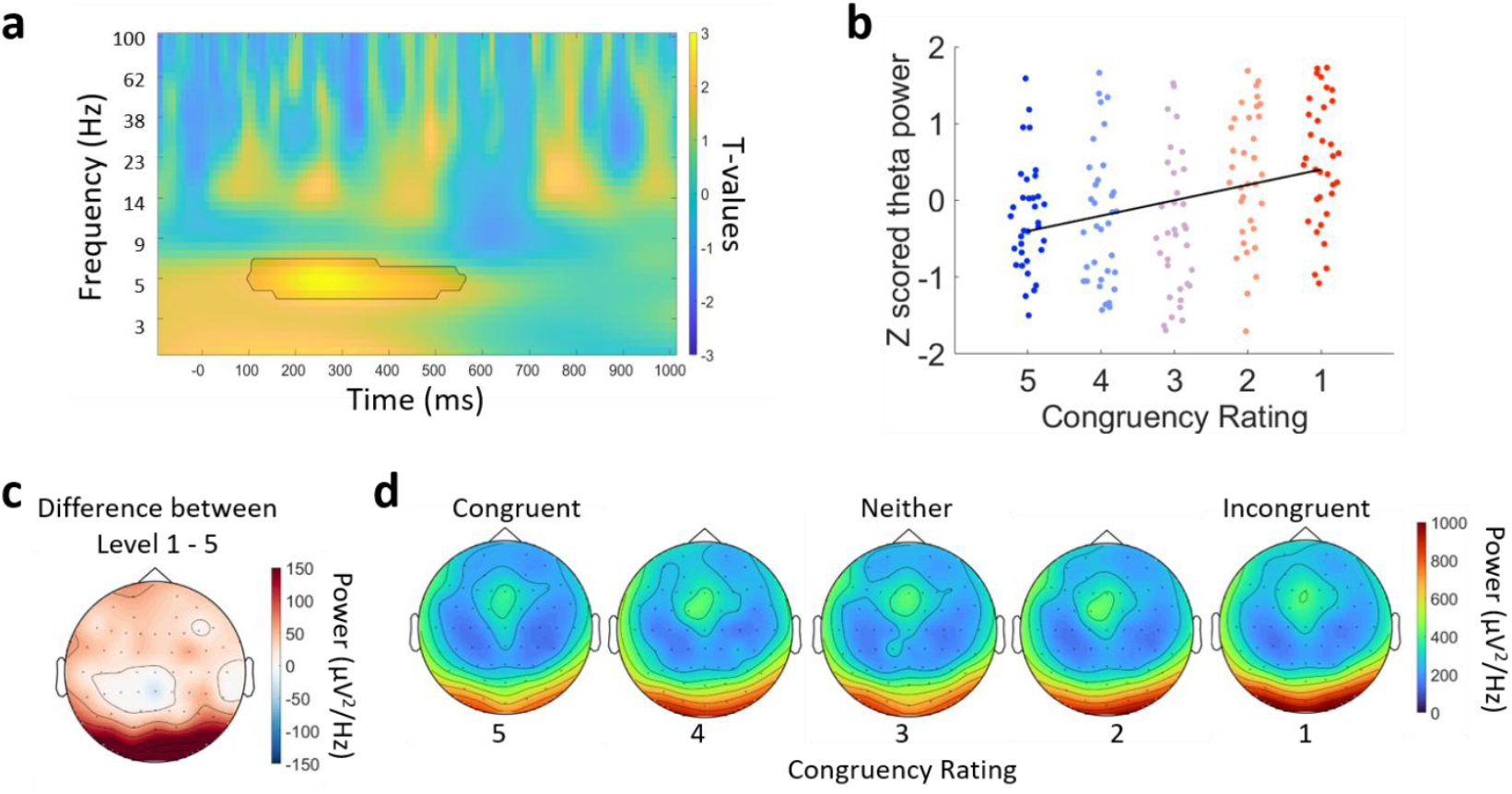
Time-frequency effects for the participant congruency ratings. (a) Time frequency representation of the contrast between incongruent and congruent trials for electrode O2 from the posterior cluster. The black line indicates the significant cluster at this electrode. (b) Z-scored power from the significant cluster for the 5 congruency levels. Each point is an individual participant. The black line is a first-degree polynomial fitted to the z-scored power. (c) Topography of the mean difference between congruency levels 1 (highly incongruent) and 5 (highly congruent) conditions averaged over 5 to 7 Hz power and between 200 and 500 ms. (d)Topographies for each of the congruency levels ranging from highly congruent on the left, to highly incongruent on the right, showing the mean power averaged over 5 to 7 Hz and between 200 and 500 ms.

## Discussion

Here we set out to determine if neural activity is impacted by context, when the context is a real spatiotemporally coherent environment, the real world. Using mEEG and AR we examined whether neural object recognition processes are impacted by real-world outdoor and indoor contexts. We found that the congruency between objects and the environment modulated both ERPs and theta-band power, with significantly greater responses for objects in incongruent contexts than objects in congruent contexts. This demonstrates that real-world contexts impact neural activity when we recognise objects, and that mobile brain imaging and AR are effective tools to study cognition in the wild.

Our main finding concerned context effects in the theta band across posterior electrodes between 100 and 600 ms, where theta-band power was higher for objects that were incongruent with the environment than for objects that were congruent with the environment. This was seen for both the predefined congruent conditions and when using the participant ratings. This is in line with previous research showing an increase in theta power when targets are incongruent, for example critical words not matching the preceding sentence (Bastiaansen, et al, 2005; Willems, Oostenveld, & Hagoort, 2008; Davidson, & Indefrey, 2007; Klimesch, 1999), which has been suggested to reflect memory processes relating to the integration of semantic information into the currently held representations of the semantic context (Bastiaansen, et al, 2005; Willems, Oostenveld, & Hagoort, 2008; Klimesch, 1999, Riddle, et al, 2020; Metzner, et al, 2015; Hald, Bastiaansen, & Hagoort 2006). Our results expand on these findings by showing that even in real spatiotemporally consistent environments, expectations about which objects appear in the environment influences how objects are recognised, and that further, this effect appears to scale with the degree of incongruence. These results give promising support to findings from laboratory studies applying to real world scenarios, even though the laboratory studies are highly reduced compared to the real world.

Theta oscillations have also been linked to the ongoing semantic processing of objects and words in the ventral visual pathway, at a similar time range to our modulations of theta activity (Clarke, Devereux, & Tyler 2018; Clarke 2020; Halgren, et al, 2015). This might suggest that the knowledge we have about the environment we are in, modulates how we access semantic information, which is primarily reflected through theta oscillations. This is also consistent with recent EEG work showing that semantic representations for objects that are incongruent with the scene are associated with enhanced and extended sematic processing compared to more congruent objects (Krugliak, et al, 2024). This points towards a system whereby the semantic processing of objects that are incongruent with the environment show enhanced theta activity as a consequence of the object recognition process not benefiting from the schema-based predictions generated by the immersion in a particular environment. This hypothesis that the increased activity for incongruent conditions compared to congruent conditions arises from the degree of expectations the environment affords is also in line with predictive coding accounts and contextual facilitation models (e.g. Eddine et al., 2024, Bar, 2004). For example, a predictive coding computational model of language comprehension can simulate many different N400 effects through summing the lexico-semantic prediction error signals, with prediction error scaling with unpredictability of a word (Eddine et al., 2024). Our congruency effects also scaled with unexpectedness of the objects in the visual environment, consistent with the models predictions, albeit in a different modality. This further suggest that the modulations we see are of a semantic nature, and reflect modulations of accessing semantic information given the environment.

While our EEG analyses do not allow us to make claims about the neural regions involved, it is noteworthy that previous studies suggest that the precuneus might integrate object sensory information with expectations generated prior to the presentation of an object (Brandman, & Peelen, 2017; Clarke, Crivelli-Decker, & Ranganath, 2022), which would correspond with the location of our theta effects over occipital and parietal electrodes. The intriguing question of how activity in this posterior medial system interacts with the ventral visual pathway could be explored in future studies using more spatially specific methods such as MEG or fMRI.

In addition to our oscillatory analysis, we also examined ERPs. Previous ERP research examining the impact context has on the neural processes involved in object recognition has found an impact on the N300 and N400 components (Coco, Aruajo, & Petersson, 2017; Coco, Nuthman, & Dimigen, 2020; Draschkow, et al, 2018), with a greater amplitude being shown for objects that are incongruent with the environment than objects that are congruent with the environment. This amplitude increase has been associated with an increase in semantic processing and enhanced lexico-semantic prediction errors. Our ERP analysis identified two clusters showing a modulation with congruency, which was restricted to our analysis of the congruency conditions. ERPs over the left anterior electrodes showed a classic N400 waveform, which was additionally modulated by congruency. The location of this effect may be different to some previous research due to differences in reference electrode, although N400 semantic effects are also commonly localized to the left anterior temporal lobes (Nobre & McCarthy, 1995; Halgren et al., 2002; Jackson et al., 2015; Lau et al., 2016), which is consistent with our ERP effects. We also saw an additional cluster over the posterior electrodes. If the ERP modulation we observe does reflect changes in the object recognition process, then the posterior and anterior clusters could reflect an increased interaction between posterior and anterior regions during recognition (Clarke, 2019). While this is likely beyond the current capabilities of mobile EEG, additional MEG or fMRI studies could shed light on this hypothesis.

Recently there has been increasing interest in examining human neural processes where participants perform tasks in more naturalistic contexts, such as VR or even the real world, whilst performing increasingly natural behaviours (Gramann, et al, 2014; Makeig, et al, 2009; Niso, et al, 2023; Iggena, et al, 2023; Gramann, et al, 2021; Hilton, Kapaj, & Fabrikant, 2024; Delaux, et al, 2021). We make a significant expansion to these studies by demonstrating the use of AR in combination with mEEG to investigate fundamental neural processes such as those underpinning object recognition. This novel combination opens the door to a whole new direction for neuroscience research, where not only can one conduct highly controlled studies in the laboratory or in highly realistic VR, but also studies in real-world locations beyond the lab. This allows for the exploration of neuroscience while participants perform natural behaviours within real environments, giving us a way to understand our neural processes involved in going about our day to day lives. Moreover, AR allows us to explore how we can change our environments and the efficiency of neural processing. For example, one could examine whether we can improve memory when viewing augmented information compared to information presented as text or static displays (e.g. Mahmoudi, Badie, & Valipour, 2015; Capuano, et al, 2016). Mobile EEG and AR also allows us to explore these questions in diverse arenas, such as museums and galleries, where mixed reality could enhance memory and visitor engagement.

Mobile imaging studies with more naturalistic designs necessarily lead to adjustments in the flow of a study. In our paradigm, participants were finding cues and walking through the environment for up to an hour. Each trial required that participants identified the physical location of the cue and travelled there, before seeing the object and providing a rating. Successive trials required participants to walk to a different location, for example a neighbouring room, meaning the pace of the experiment was slowed, and the number of trials it was possible to collect was reduced compared to a lab-based study. However, this reflects a trade-off between a more naturalistic design and one where hundreds of trials can be presented with short intervals.

One limitation of our paradigm, is that the objects were not permanently embedded into the environment, but rather appeared on a button press. We chose this as a first step towards testing object recognition in the real world with mobile EEG and AR, as it allowed us to have control over the objects and a clear time period to analyse. However, it enforces an artificial temporal constraint which would not normally be present in the real world when examining objects. As these methods and analytical approaches develop, it will become important to be able to track object recognition in a more ecologically valid way if we are to study natural cognitive processes. This may be facilitated through continued development of the integration of head-mounted AR devices, eye tracking, body sensors and mobile brain imaging, where the visual and semantic processing of our visual world can be modelled continuously depending on what we fixate upon and choose to interact with.

## Conclusion

This study provides some of the first insights into how interactions between objects and real environments modulate neural activity. Previously, research has described in detail how object congruency modulates object recognition processes in well controlled laboratory environments. However, what has been missing is information on how this process occurs when participants are present in the same environment as the stimulus objects, the environment is relatively stable and long lasting, and while participants perform naturalistic behaviours. We find that even real environments impact object recognition processes. These results are a positive for laboratory studies as it would seem even with low ecological validity, some findings appear universal, or to be similar in highly ecologically valid conditions. Moreover, we suggest mobile brain imaging and AR are excellent tools to study real world cognition.

## Data materials and availability

All data, code, and materials can be found on zenodo at: https://zenodo.org/doi/10.5281/zenodo.11394630

## Acknowledgements

This research was funded in whole, or in part, by the Wellcome Trust [Grant number 211200/Z/18/Z to AC]. For the purpose of open access, the author has applied a CC BY public copyright licence to any Author Accepted Manuscript version arising from this submission.

