## Supplemental Table 1 and Figures 1 and 2 for "Contextual Expectations in the Real-World Modulate Low-Frequency Neural Oscillations"

Victoria I. Nicholls, *et al*.

**This PDF includes:**

Supplementary Figs. 1 to 3.

Supplementary Table 1.

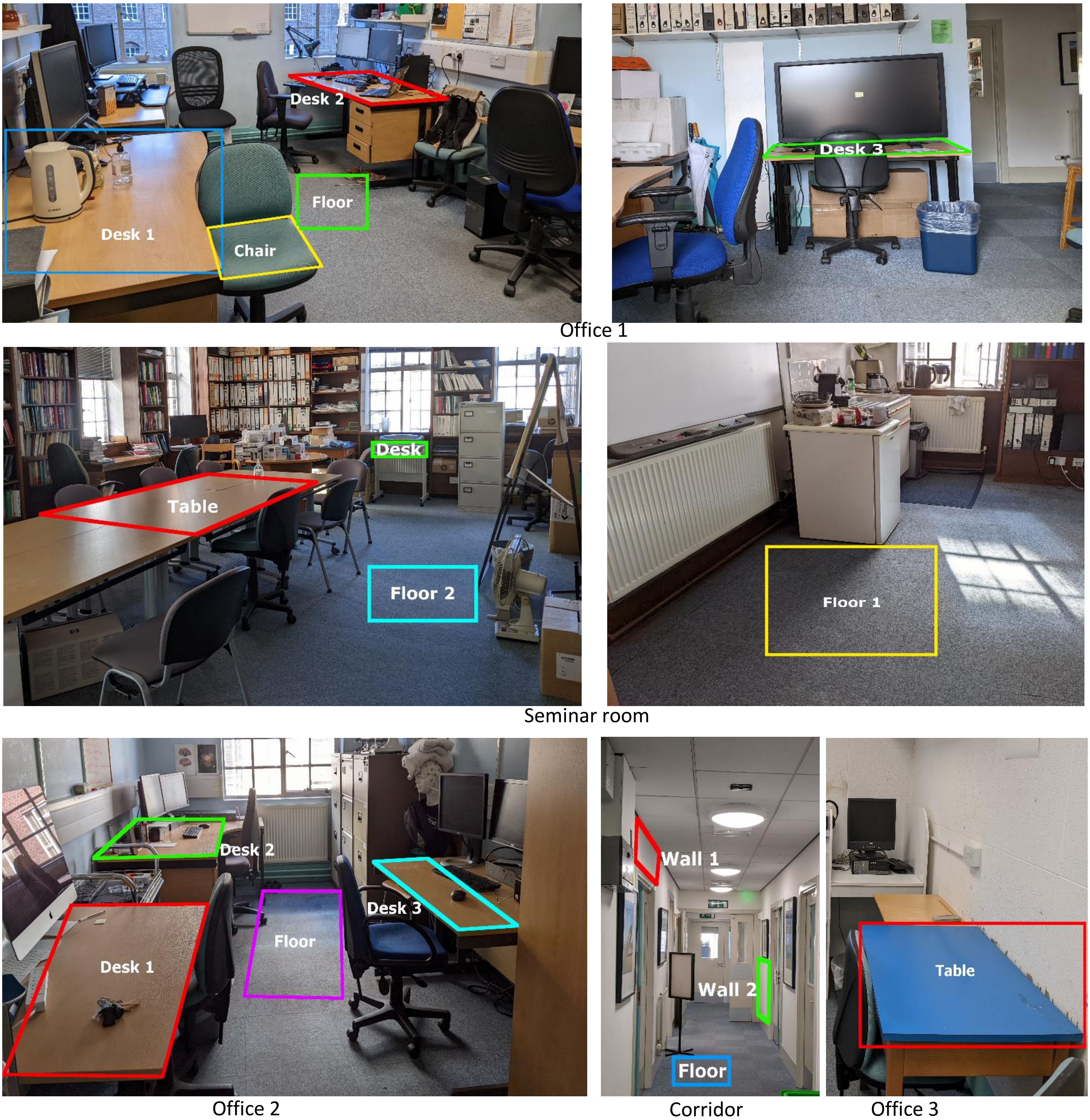


***Fig. 1 Objects and their locations in the indoor area of the experiment*** *The locations in which objects will be placed during the experiment for the indoor environment. The top row shows the locations for Office 1. The second row for the Seminar Room. The bottom row for Office 2, the corridor, and Office 3. See Supplementary Table 1 for which objects will be placed in which locations.*

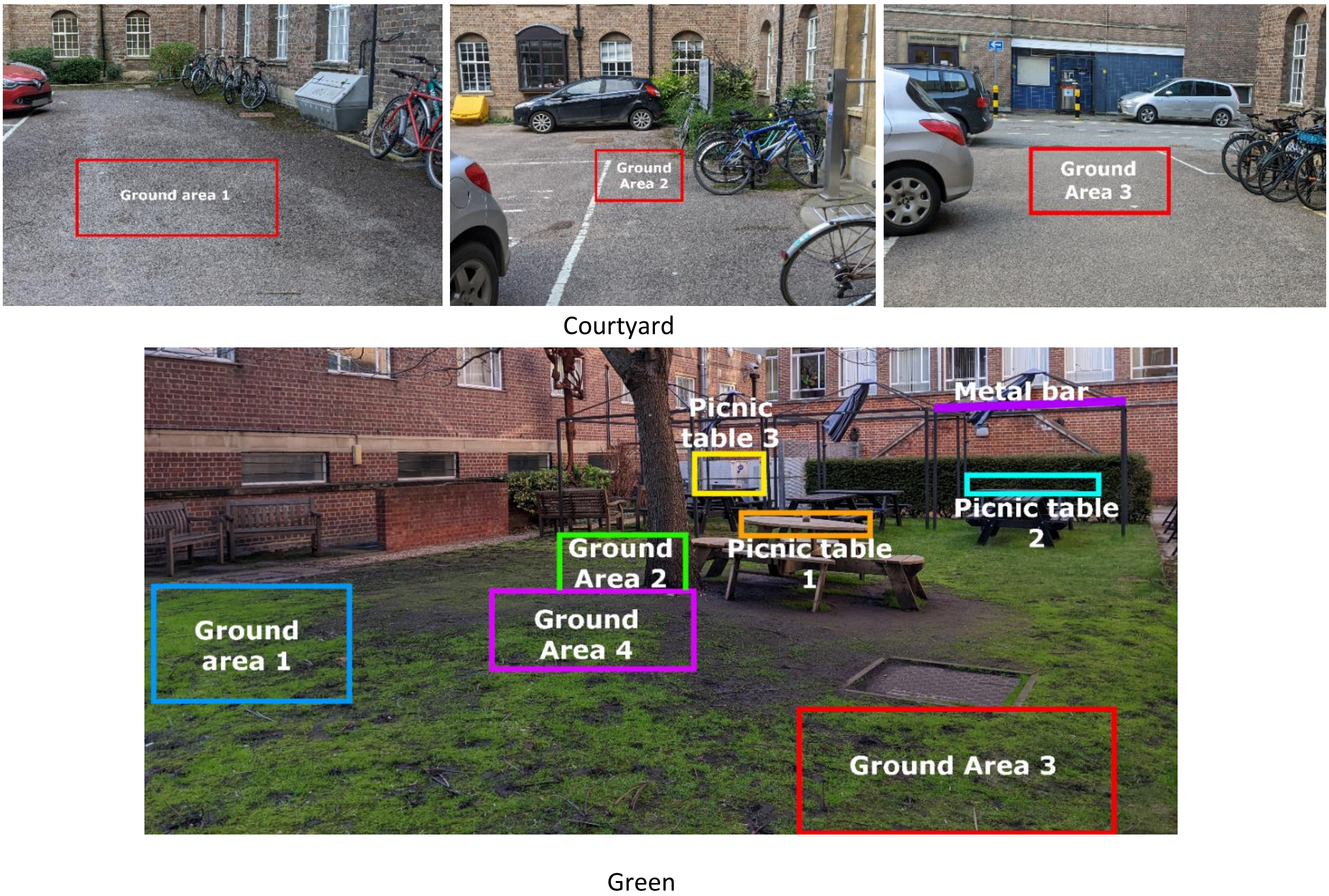


***Fig. 2 Objects and their locations in the outdoor area of the experiment.*** *The object locations for the outdoor environment. The top row shows the object locations for the Courtyard, and the bottom row for the Green. See Supplementary Table 1 for a list of which objects will be placed in which locations.*

***Table** 1 **A table of the objects that will be used in the experiment, and which environment and location the objects will be placed in**. The trial order is the order if participants perform the blocks in the order 1, 2, 3, 4. However, the blocks will be counterbalanced so participants will always perform the trials in this order. For the congruency rating 1 is congruent and 2 is incongruent with the environment.*

| Trial | Object | Environment | Location | Block | Congruency |
|---|---|---|---|---|---|
| 1 | Pen | Indoor | desk1, office 1 | 1 | 1 |
| 2 | vacuum | Indoor | floor, Office1 | 1 | 2 |
| 3 | holly | Indoor | desk1, office 2 | 1 | 2 |
| 4 | grill | Indoor | seminar room floor 1 | 1 | 2 |
| 5 | pillow | Indoor | chair, office 1 | 1 | 1 |
| 6 | mug tankard | Indoor | table, seminar room | 1 | 1 |
| 7 | monitor | Indoor | desk1, office 1 | 1 | 1 |
| 8 | binoculars | Indoor | table, seminar room | 1 | 2 |
| 9 | speaker | Indoor | table, office 3 | 1 | 1 |
| 10 | coffee mug | Indoor | desk 2, office 1 | 1 | 1 |
| 11 | hand saw | Indoor | desk 2, office 2 | 1 | 2 |
| 12 | bird house | Indoor | wall1, corridor | 1 | 2 |
| 13 | shaker | Indoor | desk3, office 1 | 1 | 2 |
| 14 | key | Indoor | table, seminar room | 1 | 1 |
| 15 | briefcase | Indoor | desk3, office 2 | 1 | 1 |
| 16 | step stool | Indoor | floor, office 2 | 1 | 2 |
| 17 | orchid | Indoor | desk1, office 2 | 1 | 1 |
| 18 | spoon | Indoor | table, seminar room | 1 | 1 |
| 19 | beer | Indoor | desk1, office 1 | 1 | 2 |
| 20 | slug | Indoor | floor, corridor | 1 | 2 |
| 21 | CD | Indoor | desk 2, office 2 | 2 | 1 |
| 22 | computer | Indoor | table, office 3 | 2 | 1 |
| 23 | moth | Indoor | wall2, corridor | 2 | 2 |
| 24 | funnel | Indoor | table, seminar room | 2 | 2 |
| 25 | office chair | Indoor | floor, office 2 | 2 | 1 |
| 26 | scotch tape | Indoor | table, seminar room | 2 | 1 |
| 27 | waffle iron | Indoor | desk 2, office 2 | 2 | 2 |
| 28 | French fries | Indoor | desk1, office 1 | 2 | 2 |
| 29 | envelopes | Indoor | desk, seminar room | 2 | 1 |
| 30 | laptop | Indoor | table, office 3 | 2 | 1 |
| 31 | headphones | Indoor | desk1, office 1 | 2 | 1 |
| 32 | toaster | Indoor | table, office 3 | 2 | 2 |
| 33 | plunger | Indoor | floor 2, seminar room | 2 | 2 |
| 34 | bowl | Indoor | table, seminar room | 2 | 1 |
| 35 | clamp | Indoor | table, office 3 | 2 | 2 |
| 36 | food processor | Indoor | desk3, office 2 | 2 | 2 |
| 37 | ruler | Indoor | desk 2, office 1 | 2 | 1 |
| 38 | bracelet | Indoor | table, seminar room | 2 | 1 |
| 39 | hairbrush | Indoor | desk1, office 1 | 2 | 1 |
| 40 | dryer | Indoor | floor 1, seminar room | 2 | 2 |
| 41 | bike helmet | Outdoor | picnic table 1, green | 3 | 1 |
| 42 | fork | Outdoor | picnic table 2, green | 3 | 1 |
| 43 | vase | Outdoor | picnic table 1, green | 3 | 2 |
| 44 | muffin | Outdoor | picnic table 3, green | 3 | 1 |
| 45 | tape | Outdoor | picnic table 1, green | 3 | 2 |
| 46 | Swiss army knife | Outdoor | picnic table 2, green | 3 | 1 |
| 47 | cat | Outdoor | ground area1, green | 3 | 1 |

| | | | | | |
|---|---|---|---|---|---|
| 48 | corkscrew | Outdoor | picnic table 2, green | 3 | 2 |
| 49 | crow | Outdoor | ground area1, green | 3 | 1 |
| 50 | grater | Outdoor | picnic table 3, green | 3 | 2 |
| 51 | squirrel | Outdoor | picnic table 1, green | 3 | 1 |
| 52 | bench | Outdoor | ground area1, green | 3 | 1 |
| 53 | sheep | Outdoor | ground area2, green | 3 | 2 |
| 54 | hanger | Outdoor | ground area 1, courtyard | 3 | 2 |
| 55 | sneakers | Outdoor | ground area 2, courtyard | 3 | 2 |
| 56 | cupboard | Outdoor | ground area 1, courtyard | 3 | 2 |
| 57 | bicycle | Outdoor | ground area 2, courtyard | 3 | 1 |
| 58 | sandal | Outdoor | ground area 1, courtyard | 3 | 2 |
| 59 | cigarette | Outdoor | ground area 2, courtyard | 3 | 1 |
| 60 | nut | Outdoor | ground area 1, courtyard | 3 | 2 |
| 61 | pine | Outdoor | picnic table 1, green | 4 | 1 |
| 62 | kingfisher | Outdoor | metal bar, green | 4 | 2 |
| 63 | cell phone | Outdoor | picnic table 2, green | 4 | 2 |
| 64 | lipstick | Outdoor | picnic table 3, green | 4 | 2 |
| 65 | dragonfly | Outdoor | picnic table 2, green | 4 | 1 |
| 66 | cabinet | Outdoor | ground area1, green | 4 | 2 |
| 67 | butterfly | Outdoor | ground area2, green | 4 | 1 |
| 68 | bush | Outdoor | ground area3, green | 4 | 1 |
| 69 | deer | Outdoor | ground area4, green | 4 | 2 |
| 70 | cap | Outdoor | picnic table 1, green | 4 | 1 |
| 71 | caterpillar | Outdoor | ground area1, green | 4 | 1 |
| 72 | thermos | Outdoor | picnic table 3, green | 4 | 1 |
| 73 | turkey | Outdoor | ground area1, green | 4 | 2 |
| 74 | pasta | Outdoor | picnic table 2, green | 4 | 2 |
| 75 | packing tape | Outdoor | picnic table 3, green | 4 | 2 |
| 76 | goose | Outdoor | ground area3, green | 4 | 2 |
| 77 | measuring tape | Outdoor | ground area 2, courtyard | 4 | 2 |
| 78 | loafer | Outdoor | ground area 1, courtyard | 4 | 2 |
| 79 | reindeer | Outdoor | ground area 3, courtyard | 4 | 2 |
| 80 | swan | Outdoor | ground area 1, courtyard | 4 | 2 |

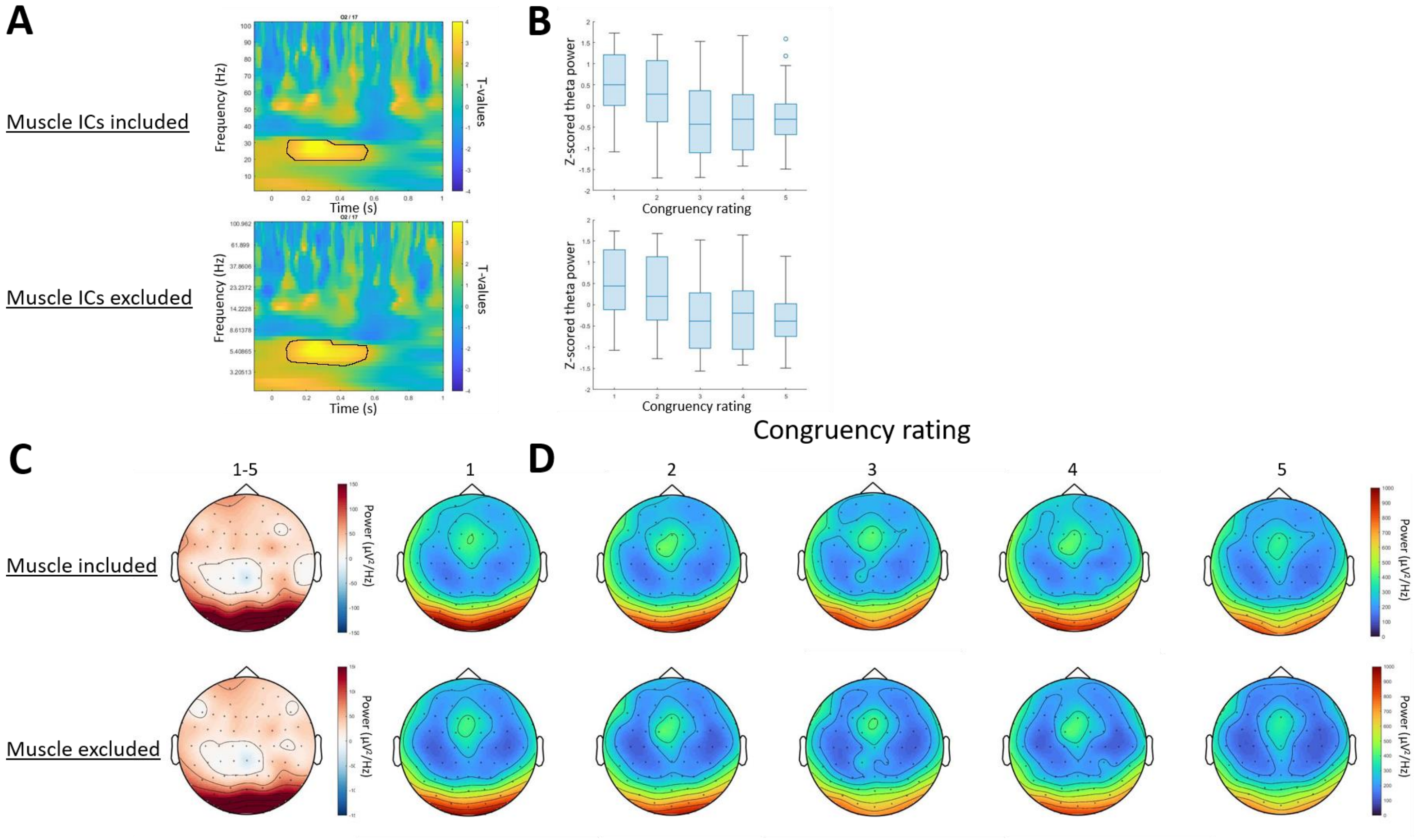


**Fig. 3. Figures demonstrating the time-frequency analyses with muscle ICs included and excluded.** (a) *T-values calculated across time and frequencies of the EEG data for electrode O2. The black line indicates the significant cluster for electrode O2. The top panel shows the results the results when muscle ICs are included and the bottom for when muscle ICs are excluded. (b) The z-scored theta power (4-7Hz) averaged across each of the five congruency ratings for each participant. The top panel shows the results the results when muscle ICs are included and the bottom for when muscle ICs are excluded. (c) The mean difference topography showing the difference in theta power between trials where objects were rated as highly incongruent (1) and highly congruent (5), between 200 and 500ms, averaged across participants. The top panel shows the results the results when muscle ICs are included and the bottom for when muscle ICs are excluded. (d) Topographies for the mean theta power (5-7Hz) between 200 and 500ms at each congruency rating sorted so that objects rated as highly incongruent (1) are on the left to objects rated as highly congruent (5) are on the right. The top panel shows the results the results when muscle ICs are included and the bottom for when muscle ICs are excluded.*
